# General assessment of genetically-encoded reporters in *Yarrowia lipolytica*

**DOI:** 10.1101/2024.12.01.625949

**Authors:** Hanqing Zhang, Luai R. Khoury, Peng Xu

## Abstract

The development of genetically-encoded reporters is important to characterize gene expression dynamics and investigate cellular-level events. Unlike Baker’s yeast, the thick cell wall and the high oil content in oleaginous yeast have restricted our ability to establish efficient fluorescence or enzyme-based reporters. In this book chapter, we detailed the protocol for how to clone and construct fluorescence reporters including hrGFP, TurboGFP, and mScarlet-I3, and the Nanoluc luciferase reporters. We quantified the fluorescence/luciferase reporter efficiency with 96-well microplate readers, flow cytometry, and confocal microscopy. Our results indicate that TurboGFP and hrGFP have a relatively low signal/noise ratio, and mScarlet-I3 yields a very high signal/noise ratio. Compared to fluorescence, luciferase Nanoluc exhibits the highest signal/noise ratio. The downside of using luciferase is the relatively laborious procedure and the related high cost. This chapter may guide us to establish an efficient and reliable reporter system to study gene expression or protein-labeling in nonconventional yeast.

## 1. Introduction

*Yarrowia lipolytica*, a generally regarded as safe (GRAS) microorganism, has become a popular and promising host for genetic and metabolic engineering because of its exceptional oleaginous ability. Such property enables this organism to produce oleochemicals [1–3], biofuels [4, 5], and acetyl CoA-derived metabolites [6, 7]. It also has been extensively studied as a model organism for lipid accumulation [8, 9] and degradation [10], dimorphism [11, 12] and protein secretory pathway [13]. In food industry, *Y. lipolytica* is also used to produce large quantities of citric acid [14, 15], α-ketoglutarate [16], and succinic acid [17]. Its advantages of utilizing various substrates including glucose, fructose, glycerol, hydrocarbon, surviving in low pH environments, make this microorganism an attractive and promising model organism for biotechnology applications. Recent notable examples of its application include the production of 2-phenylethanol [18–20], erythritol [21–23], abscisic acid [24], carotenoid [25, 26], and gastrodin [27]. Through genome sequencing [28] and tool development in recent years, the potential of *Y. lipolytica* being engineered as an industrial organism to produce novel and high-value compounds that beyond its regular portfolio of fatty acids, fatty alcohols, biofuels, and protein production is demonstrated.

Although a number of genetic tools have been established in *Y. lipolytica*, a comprehensive assessment of the reporter proteins such as various fluorescence proteins is not well-established yet. The reporter proteins are usually used for molecular imaging, such as fusion with a protein for co-localization [29], or act as a genetic marker for homologous recombination [30], etc. As for the microorganism that possesses a strong potential for industrial biotechnology like *Y. lipolytica*, reporter proteins play an extremely important role in high-throughput screening for the most efficient strain to produce high-value compounds [31, 32]. In this case, the choice of the reporter gene will be crucial for the high-throughput screening and the downstream assessment of the production of the target compounds.

In this protocol, we will describe the procedures of using different reporter genes including hrGFP [33, 34], TurboGFP [35], mScarlet-I3 [36], and nanoluciferase (Nluc) [37, 38] in *Y. lipolytica* and characterize the gene expression profile with various methods. This protocol includes the cloning of the reporter genes into the shuttle vector pYLXP’ [37, 39], and the quantification of the reporter genes in *Y. lipolytica* with microplate reader, flow cytometry, and confocal microscopy, in order to assess the protein expression level and reporter efficiency [40–43]. These characterization tools will provide a powerful platform to guide the researchers select the reporter genes wisely.

## 2. Materials

### 2.1. Reagents, media and molecular kits

1. 2 x Phanta Flash Master Mix (Dye Plus), Vazyme
2. Beyotime^®^ Seamless Cloning Kit, Beyotime Biotechnology
3. Nano-Glo® Luciferase Assay System, Promega
4. Media compositions:

1. LB medium: 16 g/L Lauria Broth powder, Agar 18 g/L (only required for agar plate), appropriate amounts of antibiotics such as 40 μg/mL kanamycin or 100 μg/mL ampicillin depending on the selective antibiotic marker on the plasmid (antibiotics are normally required except for recovery of competent cells).
2. YPD medium: 10 g/L yeast extract, 20 g/L peptone, 20 g/L glucose, 18 g/L agar (only required for agar plate).
3. CSM-Leu medium: 20 g/L glucose, 1.7 g/L yeast nitrogen base without amino acids and ammonium sulphate (YNB-AA-AS), 5 g/L ammonium sulphate, 0.7 g/L CSM-Leu powder, 18 g/L agar (only required for agar plate).

### 2.2. Strains, plasmids, and primers

Please refer to Table 1, Table 2, and Table 3 for strains, plasmids and primers for this protocol.

**Table 1.**
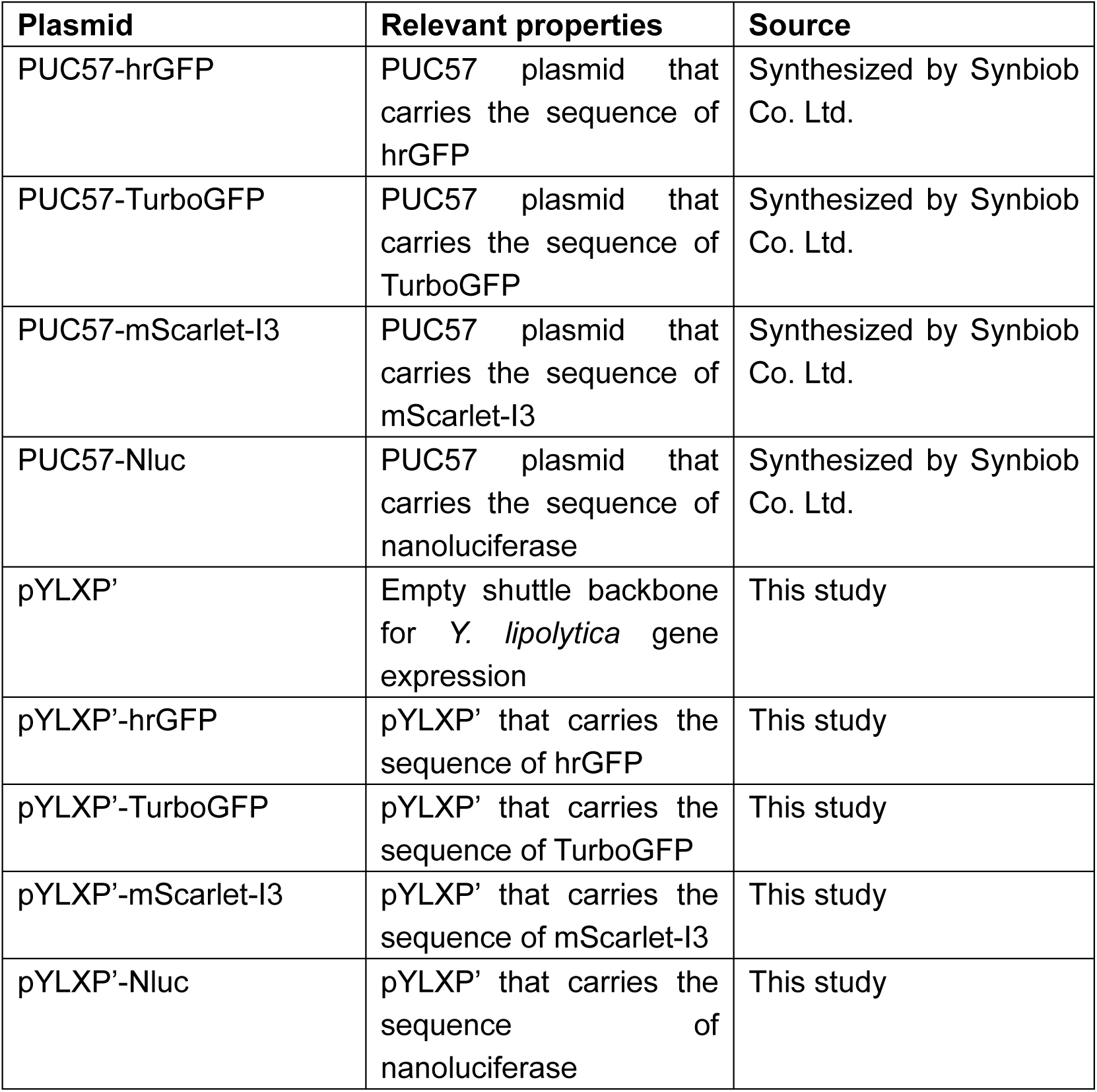
Plasmids constructed in this work.

**Table 2.** Strains.

| Strains | Genotype | Source |
| --- | --- | --- |
| *E. coli* | *fhuA2 Δ(argF-lacZ)U169 phoA glnV44 Φ80 Δ(lacZ)M15 gyrA96 recA1 relA1 endA1 thi-1 hsdR17* | Sangon Biotech |
| *Y. lipolytica* Po1g | *matA, xpr2-332, axp-2, leu2-270* | Madzak et al. |

**Table 3.**
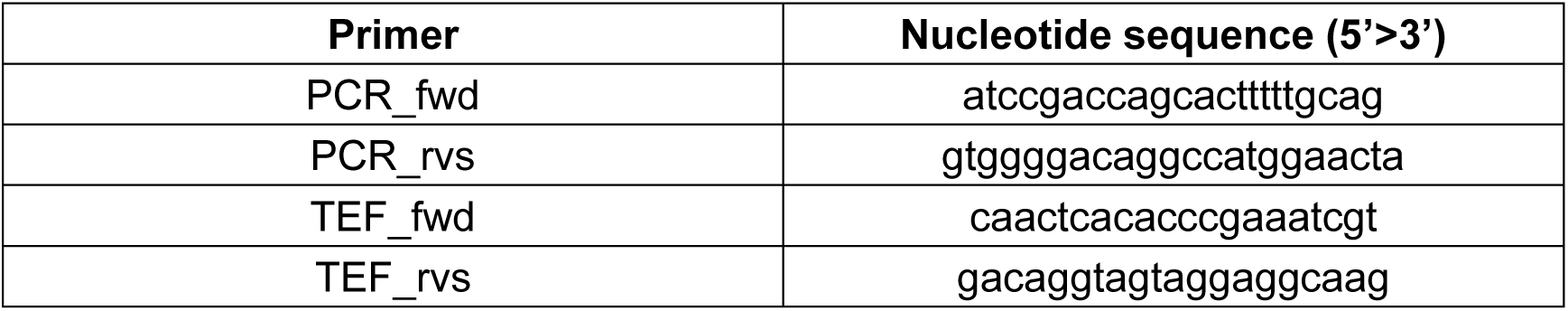
Primers.

### 2.3. Equipment

1. BioTek Synergy H1M microplate reader
2. BD LSRFortessa™ Cell Analyzer
3. Zeiss LSM 800 Confocal Laser Scanning Microscope

## 3. Methods

### 3.1. Preparation of PCR products for reporter genes

1. Inoculate *E. coli* DH5α harboring plasmid PUC57-hrGFP, PUC57-TurboGFP, PUC57-mScarlet-I3, PUC57-Nluc into 2 mL LB supplemented with 40 μg/mL of kanamycin, grow in 15 mL Corning tubes at 37 °C and 250 rpm for 16 hours in a shaker incubator.
2. Centrifuge the above cell cultures at 12,000 rpm for 1 minute to collect cell pellets.
3. Mini-preparation of the above plasmids using commercial plasmid extraction kit. Elute each plasmid with 60 μL sterilized MilliQ water.
4. Set up PCR reactions to amplify hrGFP, TurboGFP, mScarlet-I3, and Nluc gene from PUC57-hrGFP, PUC57-TurboGFP, PUC57-mScarlet-I3, PUC57-Nluc. And the following components to a PCR tube and mix:

5. Perform PCR reactions using the following parameters

| Components | Volume (for 20 μL reaction) |
| --- | --- |
| 10 μM forward primer pcr_fwd | 2 μL |
| 10 μM reverse primer pcr_rvs | 2 μL |
| Template (PUC57-hrGFP or PUC57-TurboGFP or PUC57-mScarlet-I3 or PUC57-Nluc) | 1 μL |
| 2 × Phanta Flash Master Mix (Dye Plus) | 10 μL |
| MiliQ H<sub>2</sub>O | 5 μL |

6. Prepare 0.8% agarose gel supplemented with commercial dye, after 40 minutes, load appropriate amounts of PCR products and run the gel to check for correct fragment sizes.
7. After correct verification, use 1 μL Dpn I, 2 μL 10x enzyme digestionbuffer and an appropriate volume of MiliQ water to complement each system containing different reporter PCR products to 20 μL in total. (see note 2)
8. Incubate the above systems of different reporter PCR products at 37 °C for 1 hour.
9. Perform DNA purification using the commercial PCR product purification kit to obtain purified PCR fragments for hrGFP, TurboGFP, mScarlet-I3, and Nluc.

5. Perform PCR reactions using the following parameters:
| Steps | Cycles | Temperature | Time (mm:ss) |
| --- | --- | --- | --- |
| Initial denaturation | 1 | 98 °C | 00:30 |
| Denaturation | 40 | 98 °C | 00:10 |
| Annealing |  | 58 °C | 00:05 |
| Extension |  | 72 °C | 00:05 per Kb |
| Additional extension | 1 | 72 °C | 01:00 |
| Storage | 1 | 12 °C | As needed |

### 3.2. Assembly of reporter gene onto yeast vector

1. Digest pYLXP’ with restriction enzyme SnaB I and Kpn I. For a 20 μL reaction system, use 6 μL pYLXP’ plasmid, 1 μL SnaB I, 1 μL Kpn I, 2 μL 10× enzyme digestion green buffer and 10 μL MilliQ water. Keep reactions at 37 °C for 90 minutes. At the same time, prepare 0.8% agarose gel supplemented with commercial dye. (if a enzyme digestion buffer without dye is used, remember to mix the loading dye before gel electrohphoresis so visible bands can be observed under UV light)
2. Run gel to check and purify digested pYLXP’ vectors. Load all the 20 μL digestion reaction and run gel using 1×TAE buffer at 135 V for 20 minutes. Transilluminate digested DNA fragments under UV lights and excise the gel area that contains the corresponding gene fragment. Purify the digested DNA fragments using commercial gel DNA extraction kit. Elute gene fragments with 20 μL sterilized MilliQ water. Digested gene fragments can be stored at −20 °C for future use.
3. Set up Gibson assembly reactions to clone the 4 reporter genes into the pYLXP’ vector. Incubate the reactions for 35 minutes at 50°C in a thermocycler block.

4. After the reaction completes, take the tubes out from the thermocycler for immediate use or store them at −20 °C for future use.

| Components | Volume (μL) |
| --- | --- |
| Insert | 3 (150-300ng) |
| Vector | 2 (50-150ng) |
| 2x Gibon mix (Beyotime<sup>®</sup> seamless cloning kit) | 7.5 |
| MiliQ water | Up to 15 |
| Total volume | 15 |

### 3.3. Heat shock transformation

1. Thaw chemically competent cells on ice.
2. Add up to 0.5 μg of DNA (1∼5 μL ligation reaction) to 10-50 μL DH5α Competent Cells, tap tubes gently to mix.
3. Incubate on ice for 25-30 minutes. Meanwhile, prepare a 42°C heating block with water-bath.
4. Heat shock cells for 45 seconds in a 42 °C water bath.
5. After heat shock, return tubes to the ice bath immediately and let them sit on ice for 2 minutes.
6. Add 700 μL LB medium (without antibiotics) into each tube.
7. Cap tightly and shake horizontally for 1 hour at 37 °C and 250 rpm.
8. Take out the tubes, centrifuge at 5000 rpm for 1 min, discard 600 μL supernatant.
9. Use pipette to resuspend the cells.
10. Plate all the cell cultures onto selective plates with ampicillin. While plating, make sure that the glass rod is not too hot so that it won’t kill the transformants.
11. Incubate plates for 15 hours at 37 °C.
12. Pick up colonies for colony PCR and store the plates at 4 °C. (see note 1)

### 3.4. Colony PCR verification

1. Suspend the colonies collected from above in sterilized water, use 2 μL as template, store the rest of the volumes at 4 °C.
2. Set up PCR reactions to validate the successful insertion of hrGFP, TurboGFP, mScarlet-I3, and Nluc genes from the transformants picked from different plates. Add the following components to a PCR tube and mix:

3. Perform PCR reactions using the following parameters:

| Components | Volume (for 20 μL reaction) |
| --- | --- |
| 10 μM forward primer TEF_fwd | 2 μL |
| 10 μM reverse primer XPR2_rvs | 2 μL |
| Template | 2 μL |
| 2 × Phanta Flash Master Mix (Dye Plus) | 10 μL |
| MiliQ H<sub>2</sub>O | 4 μL |

4. Prepare 0.8% agarose gel supplemented with commercial dye, after 40 minutes, load appropriate amounts of PCR products and run the gel to check for correct fragment sizes. (see note 3)
5. Take the correctly validated colonies out from 4 °C, inoculate them into 2.5 mL LB medium with 100 μg/mL ampicillin, incubate at 37 °C overnight.
6. Save 400 μL medium for cell cultures under 4 °C, centrifuge the rest of the cell cultures at 12,000 rpm for 1 minute to collect cell pellets.
7. Mini-preparation of the above plasmids using commercial plasmid extraction kit. Elute each plasmid with 60 μL sterilized MilliQ water.
8. Send the plasmids to sequencing.
9. Save the correctly sequencing-verified cell cultures with 30% sterilized glycerol solution in the volume ratio of 1:1 (e.g. Add 400 μL 30% glycerol solution into 400 μL cell culture) in −80 °C refrigerator.

3. Perform PCR reactions using the following parameters:
| Steps | Cycles | Temperature | Time (mm:ss) |
| --- | --- | --- | --- |
| Initial denaturation | 1 | 98 °C | 10:00 |
| Denaturation | 30 | 98 °C | 00:10 |
| Annealing |  | 55 °C | 00:05 |
| Extension |  | 72 °C | 00:05 per Kb |
| Additional extension | 1 | 72 °C | 01:00 |
| Storage | 1 | 12 °C | As needed |

### 3.5. *Yaorrwia lipolytica* yeast transformation

1. Prior to starting the transformation, prepare a YPD plate streaked with the appropriate host strain (in this chapter, Po1g) of *Y. lipolytica* and incubate overnight.
2. Prepare a master mix of yeast transformation buffer solution in a 200 μL PCR tube for each transformation. ssDNA should be boiled at 95°C for at least 3 minutes. The table below shows the components of one reaction.

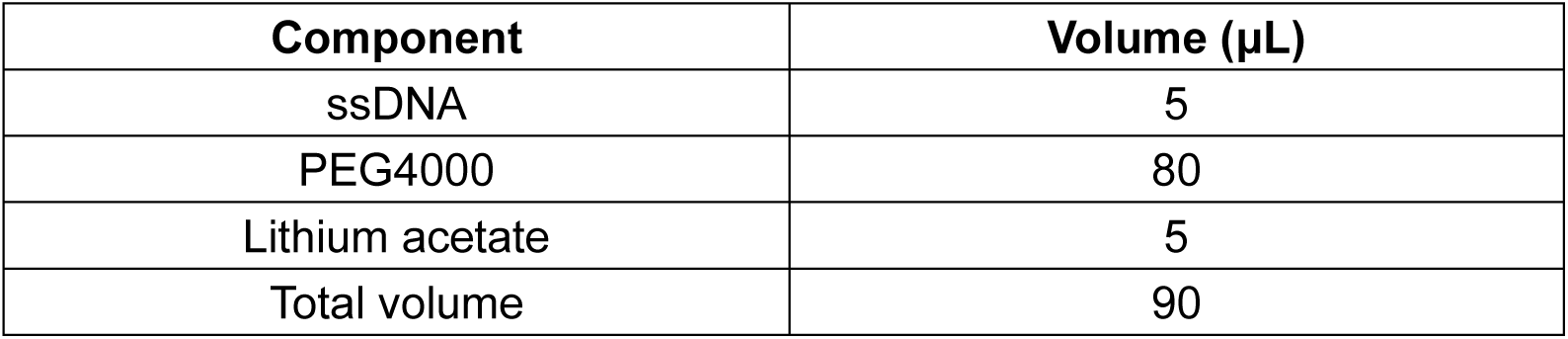
3. Scrape a lawn of *Y. lipolytic* Po1g strain from the YPD plate and transfer it to each reaction of the transformation buffer. Mix well by vortexing for about 15 seconds.
4. Add 0.5∼0.8 μg of plasmid including pYLXP’-hrGFP, pYLXP’-TurboGFP, pYLXP’-mScarlet-I3, pYLXP’-Nluc, pYLXP’, mix thoroughly by vortexing for 20 seconds.
5. Directly heat shock each transformation mixture at 39 °C and Incubate them at 39 °C for 1 hour, vortexing each transformation reaction mixture every 20 minutes for 20 seconds.
6. Dilute the cell culture by adding 100 μL of sterile water. Plate all of the diluted culture on yeast CSM-Leu plates. Incubate the plate at 30°C for at least 2 days. (see note 4)

| Component | Volume (µL) |
| --- | --- |
| ssDNA | 5 |
| PEG4000 | 80 |
| Lithium acetate | 5 |
| Total volume | 90 |

### 3.6. Characterization of hrGFP, TurboGFP, mScarlet-I3 by microplate reader

1. Prior to starting the characterization, pick three colonies from each CSM-Leu plate of four different genotypes (pYLXP’-hrGFP, pYLXP’-TurboGFP, pYLXP’-mScarlet-I3, pYLXP’), inoculate them into CSM-Leu liquid cultures at 30 °C for 36 hours.
2. Measure the value of OD600 of the above cultures, inoculate the cell cultures in a 24-well deep plate at 30 °C for 48 hours, each well contains 2 mL CSM-Leu liquid culture and the same OD600 with the value of 0.05.
3. Measure the value of OD600 of the cell cultures that are in the 24-well deep plate, dilute each cell culture to OD600 with the value of 0.2 and 0.4 with 1x PBS in a total volume of 0.5 mL.
4. Add 200 μL diluted samples from each PBS-diluted cell culture in the wells of 96-well black plate.
5. Set up the procedure of the BioTek Synergy H1 microplate reader for hrGFP and turboGFP characterization as the following table.

6. Set up the procedure of the BioTek Synergy H1 microplate reader for mScarlet-I3 characterization as the following table.

|  |  |  |
| --- | --- | --- |
| **Plate type** |  | 96 well plate |
| **Shake** | **Linear** | 0:02 (MM:SS) |
|  | **Frequency** | 567 cpm (3mm) |
| **Read: Fluorescence Endpoint** | **Excitation wavelength** | 479 nm |
|  | **Emission wavelength** | 520 nm |
|  | **Optics** | Top |
|  | **Gain** | Autoscale |
|  | **Light source** | Xenon Flash |
|  | **Lamp energy** | High |
|  | **Read speed** | Normal |
|  | **Delay** | 100 msec |
|  | **Measurements/Data points** | 10 |
|  | **Read height** | 7 mm |

7. Export and collect all the data from the software.

|  |  |  |
| --- | --- | --- |
| **Plate type** |  | 96 well plate |
| **Shake** | **Linear** | 0:02 (MM:SS) |
|  | **Frequency** | 567 cpm (3mm) |
| **Read: Fluorescence**<br>**Endpoint** | **Excitation wavelength** | 579 nm |
|  | **Emission wavelength** | 616 nm |
|  | **Optics** | Top |
|  | **Gain** | Autoscale |
|  | **Light source** | Xenon Flash |
|  | **Lamp energy** | High |
|  | **Read speed** | Normal |
|  | **Delay** | 100 msec |
|  | **Measurements/Data points** | 10 |
|  | **Read height** | 7 mm |

### 3.7. Characterization of Nluc by microplate reader

1. Prior to starting the characterization, pick three colonies from each CSM-Leu plate of two different genotypes (pYLXP’-Nluc, pYLXP’), inoculate them into CSM-Leu liquid cultures at 30 °C for 36 hours.
2. Measure the value of OD600 of the above cultures, inoculate the cell cultures in a 24-well deep plate at 30 °C for 48 hours, each well contains 2 mL CSM-Leu liquid culture and the same OD600 with the value of 0.05.
3. Measure the value of OD600 of the cell cultures that are in the 24-well deep plate, dilute each cell culture to OD600 with the value of 0.2 and 0.4 with 1x PBS in a total volume of 0.5 mL.
4. To perform the luciferase assay, equilibrate the Nano-Glo® Luciferase Assay Buffer to room temperature.
5. Each reaction in a 384-well white plate requires the following (It is recommended to make a master mix of the assay buffer and substrate first. Aliquot the lysates into the wells. Try to separate each test well with one blank well to avoid the interference of the neighbor well):

6. Set up the procedure of the BioTek Synergy H1 microplate reader for Nluc characterization as the following table.

| **Component** | **Volume (μL)** |
| --- | --- |
| Nano-Glo® Luciferase Assay Buffer | 30 |
| Nano-Glo® Luciferase Assay Substrate | 0.6 |
| Cell culture | 30 |
| Total | 60.6 |

7. Export and collect all the data from the software.

6. Set up the procedure of the BioTek Synergy H1 microplate reader for Nluc characterization as the following table.
|  |  |  |
| --- | --- | --- |
| **Plate type** |  | 384 well plate |
| **Shake** | **Linear** | 0:02 (MM:SS) |
|  | **Frequency** | 567 cpm (3mm) |
| **Start kinetics** | **Runtime** | 0:30:00 (HH:MM:SS) |
|  | **Interval** | 0:00:56 |
|  | **Reads** | 33 |
| **Read: Luminescence Endpoint** | **Integration time** | 0:01.00 (MM:SS:ss) |
|  | **Emission** | Full light |
|  | **Optics** | Top |
|  | **Gain** | Autoscale (see note 5) |
|  | **Read speed** | Normal |
|  | **Delay** | 100 msec |
|  | **Read height** | 1 mm |

### 3.8. Characterization of hrGFP, TurboGFP, mScarlet-I3 by flow cytometry

1. Prior to starting the characterization, pick three colonies from each CSM-Leu plate of four different genotypes (pYLXP’-hrGFP, pYLXP’-TurboGFP, pYLXP’-mScarlet-I3, pYLXP’), inoculate them into CSM-Leu liquid cultures at 30 °C for 36 hours.
2. Measure the value of OD600 of the above cultures, inoculate the cell cultures in a 24-well deep plate at 30 °C for 48 hours, each well contains 2 mL CSM-Leu liquid culture and the same OD600 with the value of 0.05.
3. Extract 1 mL cell culture from each well, centrifuge the extracted cell cultures at 2800 rpm for 5 minutes, discard the supernatants, add 1 mL 1x PBS buffer into each tube, and resuspend the cell pellets.
4. Measure the value of OD600 of PBS-washed cultures, dilute each sample to an OD600 value of 0.01 with 1x PBS buffer. (see note 6)
5. Start the opening procedure of flow cytometry following the corresponding instructions of the equipment.
6. Load the negative control (i.e. the genotype of pYLXP’) first.
7. Set the number of “Stopping Cells Events” to 10000, start acquiring the data.
8. Obtain the graph of FITC-A channel vs. counts, adjust the PMT voltage until the spots basically all fall below the FITC-A value of 10^2^ in log scale, keep the PMT voltage the same during all the characterization. Click “Record Data” to obtain the data of hrGFP, TurboGFP, and mScarlet-I3.
9. Load other samples, acquiring the data for each sample, and click “Record Data” for each sample.
10. After completing all the experiment, enter shut down procedure following the instruction of the equipment.
11. Collect all the data from the software.

### 3.9. Characterization of hrGFP, TurboGFP, mScarlet-I3 by confocal microscope

1. Prior to starting the characterization, pick three colonies from each CSM-Leu plate of four different genotypes (pYLXP’-hrGFP, pYLXP’-TurboGFP, pYLXP’-mScarlet-I3, pYLXP’), inoculate them into CSM-Leu liquid cultures at 30 °C for 36 hours.
2. Measure the value of OD600 of the above cultures, inoculate the cell cultures in a 24-well deep plate at 30 °C for 48 hours, each well contains 2 mL CSM-Leu liquid culture and the same OD600 with the value of 0.05.
3. Extract 1 mL cell culture from each well, centrifuge the extracted cell cultures at 2800 rpm for 5 minutes, discard the supernatants, add 1 mL 1x PBS buffer into each tube, and resuspend the cell pellets.
4. Measure the value of OD600 of PBS-washed cultures, dilute each sample to an OD600 value of 0.4 with 1x PBS buffer.
5. Start the equipment following the guide.
6. Drip 10 μL from one sample on a positively charged surface of the glass slide, carefully cover with a cover glass.
7. Select the 40x or 63x objective lens, drip a drop of Carl Zeiss™ Immersol™ Immersion Oil on the chosen lens, place the glass slide on the lens with the cover glass facing down.
8. Observe the sample in bright field to find an appropriate area and adjust the focal length.
9. Switch to Acquisition mode, set up the excitation wavelength of 479 nm and emission wavelength of 520 nm as the channel for hrGFP and TurboGFP, set up the excitation wavelength of 579 nm and emission wavelength of 616 nm as the channel for mScarlet-I3.
10. Select the desirable channels including bright field.
11. Acquire the image in live mode to find a suitable sample area in bright field, switch to the channel of hrGFP and TurboGFP to observe the fluorescence of hrGFP and TurboGFP or the channel of mScarlet-I3 to observe the fluorescence of mScarlet-I3.
12. Adjust the Master Gain of the image up to 900 V to modulate the fluorescence signal. (see note 7)
13. Click “Snap” to obtain the image.
14. After finishing all the sample characterization, shut down the confocal microscope following the instructions of the equipment.
15. Collect all the data from the software.

## 4. Notes

1. If colonies are not formed after Gibson assembly cloning, try to adjust the incubation time of the 50 °C assembly reaction, and follow the molar ratio of the vector and insert suggested by the instruction in the protocol of the Gibson assembly kit.
2. Try to do gel purification if there are non-target bands after PCR amplification of target gene fragments when running gel electrophoresis. But if the target band is much brighter than the non-target bands, it is acceptable to proceed to the PCR product purification.
3. After the gel electrophoresis of colony PCR, if false positive rates are too high to find the correct verified colony, try to prolong the incubation time of plasmid backbone digestion, or check the primers and run another PCR again with higher annealing temperature and do gel purification to purify the target PCR fragments.
4. The yeast transformation can be considered a failure if no colonies grow on the plate after incubation for three days. Try to do the transformation again with fresh Po1g colonies, make sure to add correct volumes of the transformation components, and mix well.
5. The Gain value for characterization by microplate reader can be selected by someone’s interest or Autoscale option, the key is to keep the Gain value the same among the characterization of the same reporter.
6. The OD600 value of characterization by flow cytometry can also be adjusted by researchers. A low OD600 value can prevent the clogging of the flow cytometer, so it is better to keep a low OD600 value for testing if there is no other requirement for an OD600 value.
7. Confocal microscopy for *Y. lipolytica* reporter characterization is troublesome and sometimes may be less accurate since the yeast can also generate the fluorescence signal with a similar wavelength as GFPs and mScarlet-I3 because of the lipid bodies inside the yeast. When observing the reporter performance using a confocal microscope, it is necessary to adjust the Master Gain of the figure to eliminate the background noise until some significant bright signals are trapped in some cells.
8. By comparison of the signals of different reporters using microplate reader, Nluc has extremely high fold-change signals that far exceed the fold-change signals of the other three fluorescent proteins for both OD=0.2 and OD=0.4 (Figure 1). However, it is not suitable for flow cytometry and confocal microscope if the cooled charge-coupled device (CCD) is not available for single-cell level evaluation.

**Figure 1.**
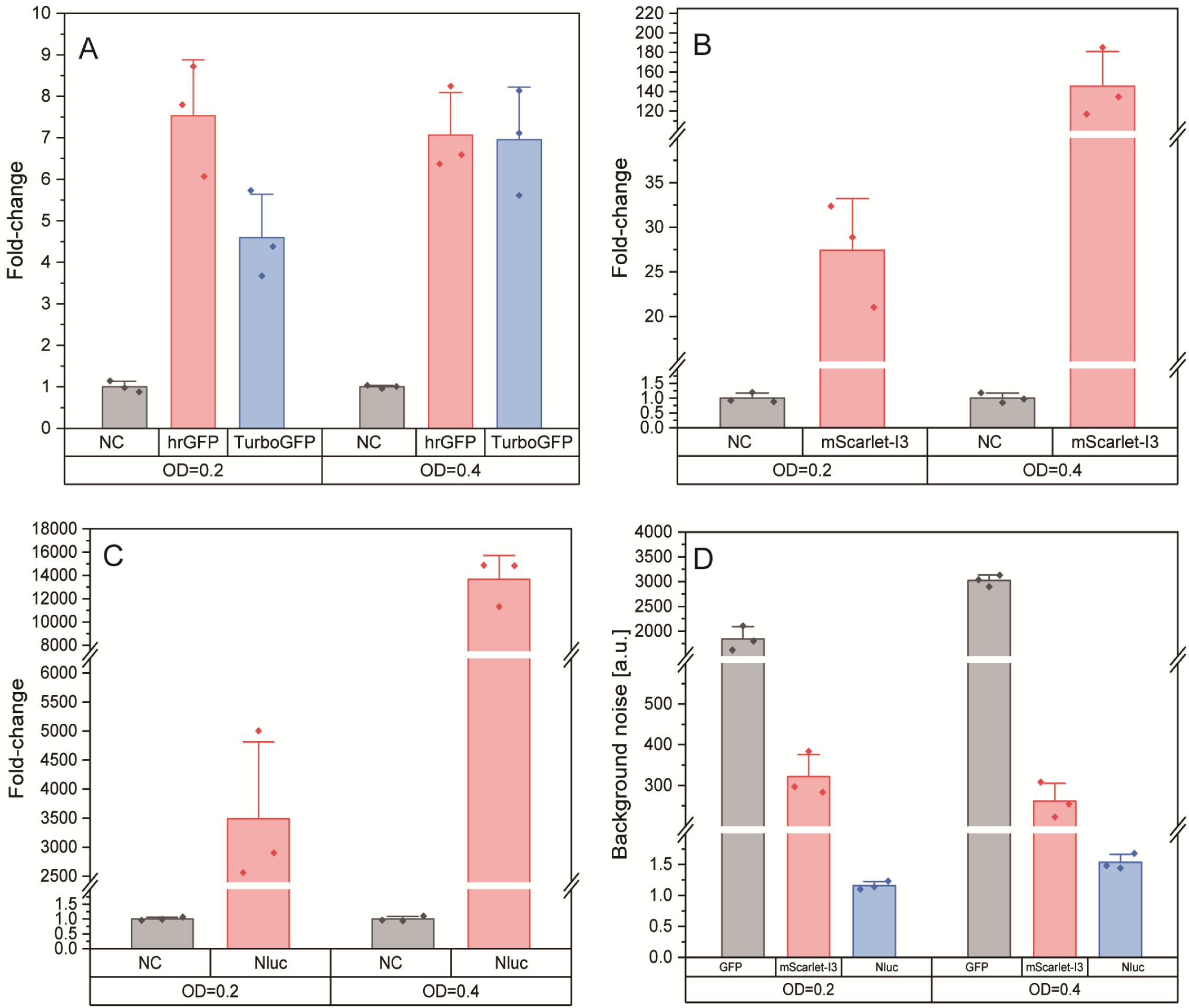
Characterization of hrGFP, TurboGFP, mScarlet-I3, and Nluc in *Y. Lipolytica* using the microplate reader at different values of OD600. A) Analysis for hrGFP and TurboGFP. B) Analysis for mScarlet-I3. C) Analysis for Nluc. D) Background noise analysis for the three reporters and the different OD600 values. NC, negative control. The error bars represent the standard deviation of n=3 biologically independent experiments. (see note 8, 9, 10, 14, 15)
9. Among the three fluorescent proteins, mScarlet-I3 generates much higer signals than the two green fluorescent proteins, while hrGFP is more sensitive at OD=0.2 compared to TurboGFP but shows similar signal strength when OD=0.4 (Figure 1).
10. It can be concluded from the characterization by the microplate reader that *Y. lipolytica* has nearly zero background noise of Nluc, both hrGFP and TurboGFP generate significant background noise. The background noise of mScarlet is nearly 10-fold lower than that of GFP in *Y. lipolytica*, but still very high compared to that of Nluc. This is due to the interference of the lipid bodies which also emits green fluorescence in the yeast.
11. When using flow cytometry to analyze the three fluorescent proteins, it is found that a larger population of cells can emit mScarlet-I3 fluorescence, compared to that only a small fraction of cells emits green fluorescence. And the total fluorescence for 10,000 cells also indicates the stronger singal observed in mScarlet than that of both hrGFP and TurboGFP reporters (Figure 2C-D).

**Figure 2.**
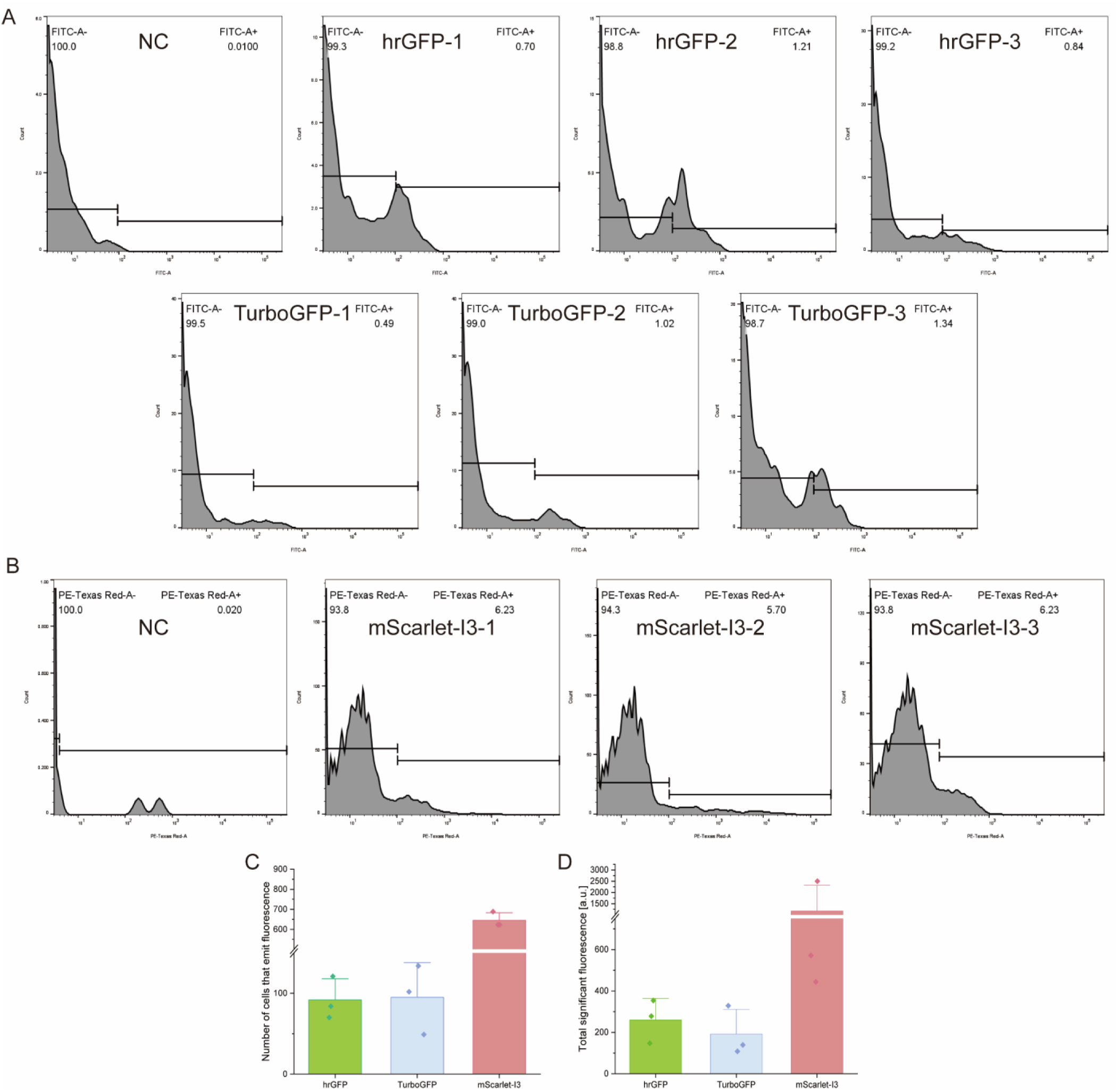
Characterization of hrGFP, TurboGFP, and mScarlet-I3 in *Y. Lipolytica* by flow cytometry. A) Results for hrGFP and TurboGFP characterization. B) Results for mScarlet-I3 characterization. C) Number of cells that have significant fluorescence signal. The numbers inside the graphs in A) and B) (e.g. 100, 0.01, 0.7, etc.) represent the percentages of the cells with the fluorescence signal within a certain range. D) Total fluorescence intensity of the cells that generate significant signals. NC, negative control. The error bars represent the standard deviation of n=3 biologically independent experiments. (see note 11, 12, 14, 16)
12. By comparing the two GFP reporters, we conclude that the total fluorescence signals for 10,000 cells are closed, suggesting that when the tested cell population is too low, there is no obvious difference between the TurboGFP and hrGFP (Figure 2C-D).
13. Similar conclusion can also be reached when comparing the image taken from confocal microscope. The cells transformed with plasmid carrying mScarlet-I3 generate stronger signals than that of the cells transformed with plasmid carrying with hrGFP or TurboGFP (Figure 3A-C).
14. mScarlet-I3 yields much stronger signal than that of two other fluorescence reporters, possibly due to the rapid maturation, high quantum yield, and intrinsic brightness of mScarlet-I3 [36].

**Figure 3.**
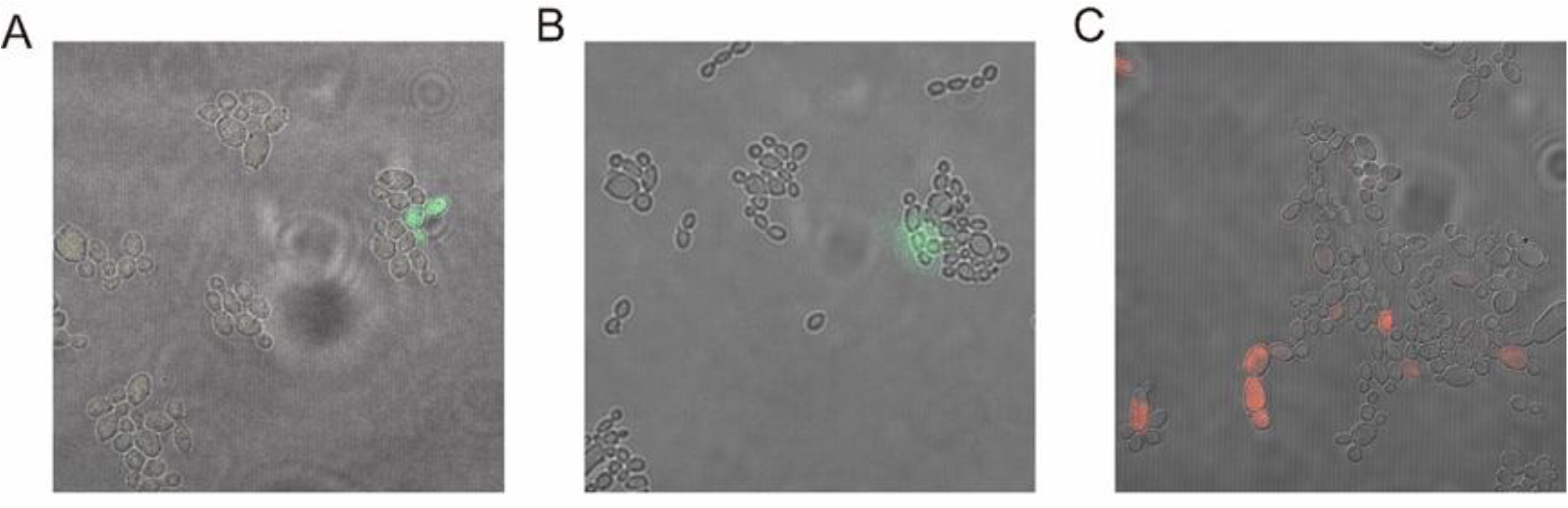
Figures taken from confocal microscope. A) hrGFP. B) TurboGFP. C) mScarlet-I3. (see note 13, 14, 16, 17)
15. Reporter proteins have various applications for *Y. lipolytica*, Nluc will be an efficient and reliable reporter if only the macroscopic evaluation is required, for example, assessing genetic tools such as CRISPRi or CRISPRa, or acting as a biosensor for high-throughput screening for the highly-productive strain.
16. Though mScarlet-I3 doesn’t emit the signal as strong as Nluc, it can serve as a single-cell level characterization tool, a good balance of signal/noise ratio and signal intensity.
17. As the two GFP reporters, they are not comparable to either Nluc or mScarlet-I3. The application of GFP in Y*. lipolytica* should best be done with confocal microscopy at single cell level.
18. This chapter would guide researchers to select the appropriate reporters in the tool developments or industrial applications in *Yarrowia lipolytica*.

## Acknowledgement

This work was supported by the National Natural Science Foundation of China (no. 22378083) and the Guangdong Provincial Key Laboratory of Materials and Technologies of Energy Conversion (MATEC) (no. GR2300014 and no. KD2200116).

## References

1. Xu, P., Qiao, K., Ahn, W. S., et al., Engineering Yarrowia lipolytica as a platform for synthesis of drop-in transportation fuels and oleochemicals. Proceedings of the National Academy of Sciences, 2016. 113(39): p. 10848–10853.

2. Abghari, A. and S. Chen, Yarrowia lipolytica as an oleaginous cell factory platform for the production of fatty acid-based biofuel and bioproducts. Frontiers in Energy Research, 2014. 2.

3. Ma, J., Gu, Y., Marsafari, M., et al., Synthetic biology, systems biology, and metabolic engineering of Yarrowia lipolytica toward a sustainable biorefinery platform. Journal of Industrial Microbiology and Biotechnology, 2020. 47(9-10): p. 845–862.

4. Ledesma-Amaro, R. and J.-M. Nicaud, Yarrowia lipolytica as a biotechnological chassis to produce usual and unusual fatty acids. Progress in lipid research, 2016. 61: p. 40–50.

5. Xu, P., K. Qiao, and G. Stephanopoulos, Engineering oxidative stress defense pathways to build a robust lipid production platform in Yarrowia lipolytica. Biotechnol Bioeng, 2017. 114(7): p. 1521–1530.

6. Zhu, Q. and E.N. Jackson, Metabolic engineering of Yarrowia lipolytica for industrial applications. Current opinion in biotechnology, 2015. 36: p. 65–72.

7. Ledesma-Amaro, R., Lazar, Z., Rakicka, M., et al., Metabolic engineering of Yarrowia lipolytica to produce chemicals and fuels from xylose. Metabolic engineering, 2016. 38: p. 115–124.

8. Beopoulos, A., T. Chardot, and J.-M. Nicaud, Yarrowia lipolytica: A model and a tool to understand the mechanisms implicated in lipid accumulation. Biochimie, 2009. 91(6): p. 692–696.

9. Qiao, K., Wasylenko, T.M., Zhou, K., et al., Lipid production in Yarrowia lipolytica is maximized by engineering cytosolic redox metabolism. Nat Biotechnol, 2017. 35(2): p. 173–177.

10. Fickers, P., Benetti, P.H., Waché, Y., et al., Hydrophobic substrate utilisation by the yeast Yarrowia lipolytica, and its potential applications. FEMS yeast research, 2005. 5(6-7): p. 527–543.

11. Ruiz-Herrera, J. and R. Sentandreu, Different effectors of dimorphism in Yarrowia lipolytica. Archives of Microbiology, 2002. 178: p. 477–483.

12. Morales-Vargas, A.T., A. Domínguez, and J. Ruiz-Herrera, Identification of dimorphism-involved genes of Yarrowia lipolytica by means of microarray analysis. Research in microbiology, 2012. 163(5): p. 378–387.

13. Beckerich, J.-M., A. Boisramé-Baudevin, and C. Gaillardin, Yarrowia lipolytica: a model organism for protein secretion studies. Int Microbiol, 1998. 1(2): p. 123–30.

14. Papanikolaou, S., Muniglia, L., Chevalot, I., et al., Yarrowia lipolytica as a potential producer of citric acid from raw glycerol. Journal of applied microbiology, 2002. 92(4): p. 737–744.

15. Papanikolaou, S., Galiotou-Panayotou, M., Fakas, S., et al., Citric acid production by Yarrowia lipolytica cultivated on olive-mill wastewater-based media. Bioresource Technology, 2008. 99(7): p. 2419–2428.

16. Zhou, J., Zhou, H., Du, G., et al., Screening of a thiamine-auxotrophic yeast for α-ketoglutaric acid overproduction. Letters in applied microbiology, 2010. 51(3): p. 264–271.

17. Cui, Z., Gao, C., Li, J., et al., Engineering of unconventional yeast Yarrowia lipolytica for efficient succinic acid production from glycerol at low pH. Metabolic engineering, 2017. 42: p. 126–133.

18. Mitri, S., Louka, N., Rossignol, T., et al., Bioproduction of 2-Phenylethanol by Yarrowia lipolytica on Sugar Beet Molasses as a Low-Cost Substrate. Fermentation, 2024. 10(6): p. 290.

19. Gu, Y., Ma, J., Zhu, Y., et al., Refactoring Ehrlich Pathway for High-Yield 2-Phenylethanol Production in Yarrowia lipolytica. ACS Synthetic Biology, 2020. 9(3): p. 623–633.

20. Gu, Y., Ma, J., Zhu, Y., et al., Engineering Yarrowia lipolytica as a Chassis for De Novo Synthesis of Five Aromatic-Derived Natural Products and Chemicals. ACS Synthetic Biology, 2020. 9(8): p. 2096–2106.

21. Rywińska, A., Tomaszewska-Hetman, L., Juszczyk, P., et al., Enhanced Production of Erythritol from Glucose by the Newly Obtained UV Mutant Yarrowia lipolytica K1UV15. Molecules, 2024. 29(10): p. 2187.

22. Qiu, X., Xu, P., Zhao, X., et al., Combining genetically-encoded biosensors with high throughput strain screening to maximize erythritol production in Yarrowia lipolytica. Metabolic Engineering, 2020. 60: p. 66–76.

23. Qiu, X., Gu, Y., Du, G., et al., Conferring thermotolerant phenotype to wild-type Yarrowia lipolytica improves cell growth and erythritol production. Biotechnol Bioeng, 2021. 118(8): p. 3117–3127.

24. Liu, L., K. Zhao, and Z. Liu, Construction and Regulation of the Abscisic Acid Biosynthesis Pathway in Yarrowia lipolytica. Journal of Agricultural and Food Chemistry, 2024. 72(13): p. 7299–7307.

25. Naveira-Pazos, C., Veiga, M.C., Mussagy, C.U., et al., Carotenoids production and extraction from Yarrowia lipolytica cells: A biocompatible approach using biosolvents. Separation and Purification Technology, 2024. 343: p. 127136.

26. Ma, Y., Liu, N., Greisen, P., et al., Removal of lycopene substrate inhibition enables high carotenoid productivity in Yarrowia lipolytica. Nature Communications, 2022. 13(1): p. 572.

27. Gu, Y., Jiang, Y., Li, C., et al., High titer production of gastrodin enabled by systematic refactoring of yeast genome and an antisense-transcriptional regulation toolkit. Metabolic Engineering, 2024.

28. Liu, L. and H.S. Alper, Draft genome sequence of the oleaginous yeast Yarrowia lipolytica PO1f, a commonly used metabolic engineering host. Genome announcements, 2014. 2(4): p. 10.1128/genomea.00652-14.

29. Miyashita, T., Confocal microscopy for intracellular co-localization of proteins. Protein-Protein Interactions: Methods and Applications, 2004: p. 399–409.

30. Vriend, L.E., M. Jasin, and P.M. Krawczyk, Assaying break and nick-induced homologous recombination in mammalian cells using the DR-GFP reporter and Cas9 nucleases, in Methods in enzymology. 2014, Elsevier. p. 175–191.

31. Groenewald, M., Boekhout, T., Neuvéglise, C., et al., Yarrowia lipolytica: safety assessment of an oleaginous yeast with a great industrial potential. Critical reviews in microbiology, 2014. 40(3): p. 187–206.

32. Lv, Y., Gu, Y., Xu, J., et al., Coupling metabolic addiction with negative autoregulation to improve strain stability and pathway yield. Metabolic Engineering, 2020. 61: p. 79–88.

33. Blazeck, J., Liu, L., Redden, H., et al., Tuning gene expression in Yarrowia lipolytica by a hybrid promoter approach. Applied and environmental microbiology, 2011. 77(22): p. 7905–7914.

34. Jin, C., Lu, Y., Jelinek, J., et al., TET1 is a maintenance DNA demethylase that prevents methylation spreading in differentiated cells. Nucleic acids research, 2014. 42(11): p. 6956–6971.

35. Maselko, M., Heinsch, S.C., Chacón, J.M., et al., Engineering species-like barriers to sexual reproduction. Nature Communications, 2017. 8(1): p. 883.

36. Gadella Jr, T.W., van Weeren, L., Stouthamer, J., et al., mScarlet3: a brilliant and fast-maturing red fluorescent protein. Nature methods, 2023. 20(4): p. 541–545.

37. Wong, L., Engel, J., Jin, E., et al., YaliBricks, a versatile genetic toolkit for streamlined and rapid pathway engineering in Yarrowia lipolytica. Metabolic engineering communications, 2017. 5: p. 68–77.

38. Masser, A.E., Kandasamy, G., Kaimal, J.M., et al., Luciferase NanoLuc as a reporter for gene expression and protein levels in Saccharomyces cerevisiae. Yeast, 2016. 33(5): p. 191–200.

39. Wong, L., Holdridge, B., Engel, J., et al., Genetic Tools for Streamlined and Accelerated Pathway Engineering in Yarrowia lipolytica, in Microbial Metabolic Engineering: Methods and Protocols, C.N.S. Santos and P.K. Ajikumar, Editors. 2019, Springer New York: New York, NY. p. 155–177.

40. Wei, T. and H. Dai, Quantification of GFP signals by fluorescent microscopy and flow cytometry. Yeast Protocols, 2014: p. 23–31.

41. Thomas, S., N.D. Maynard, and J. Gill, DNA library construction using Gibson Assembly®. Nature Methods, 2015. 12(11): p. i–ii.

42. Russell, D.W. and J. Sambrook, Molecular cloning: a laboratory manual. Vol. 1. 2001: Cold Spring Harbor Laboratory Cold Spring Harbor, NY.

43. Gietz, R.D. and R.A. Woods, Transformation of yeast by lithium acetate/single-stranded carrier DNA/polyethylene glycol method, in Methods in enzymology. 2002, Elsevier. p. 87–96.

